# Setting Starting Level for a Trial of an Biofilm-Disrupting Adjuvant

**DOI:** 10.1101/447607

**Authors:** Braden Adams, Robert A. Lodder

**Affiliations:** Department of Pharmaceutical Sciences College of Pharmacy University of Kentucky Lexington, KY 40536

## Abstract

Combination therapy is beneficial treatment modality for multiple diseases. Cinnamon oil may be an advantageous agent in a number of combination therapies as cinnamon oil has antibacterial activity. Cinnamon oil has also been shown to be effective against biofilm cultures of *Streptococcus mutans* and *Lactobacillus plantarum*. *Cinnamomum osmophloeum* inhibits planktonic cultures of many gram-positive and gram-negative bacteria, including MRSA (methicillin-resistant *Staphylococcus aureus*). Subjecting *S. epidermidis* to cinnamon oil eliminates planktonic cells or staphylococci in biofilms. Cinnamon oil is an essential oil used throughout the food industry because of its pleasant and distinctive aroma. The purpose of this study is to estimate the amount of exposure to cinnamon through selected foods in the United States in order to propose an initial target level for pharmacokinetic studies of a novel combination drug.

## Introduction

Combination therapy is an important treatment approach for many diseases, such as cancer, cardiovascular disease, and infectious diseases. A steady stream of scientific advances have improved our insight into the pathophysiological processes that form the basis of these and other complex diseases. This improved knowledge has provided new momentum for novel therapeutic approaches using combinations of drugs aimed at multiple therapeutic targets to enhance treatment response or reduce generation of resistance. In environments in which combination therapy offers important therapeutic superiority, there is increasing enthusiasm for the expansion of combinations of investigational drugs not already developed (FDA, 2010).

Cinnamon oil may be a useful agent in some combination therapies. Cinnamon oil is an essential oil widely employed in the food industry because of its unique aroma (Nuryastuti, 2009). *Cinnamomum* is a genus in the family *Lauraceae*, and many species of Lauraceae are utilized as spices. The cinnamon stick found in grocery stores in the United States is *Cinnamomum burmannii* from Indonesia, also known as Indonesian cassia. Cinnamon oil can be obtained from the bark of Indonesian cassia.

A number of papers have shown that cinnamon oil has antibacterial activity. Cinnamon oil has also been demonstrated to be active against biofilm cultures of *Streptococcus mutans* and *Lactobacillus plantarum*. *Cinnamomum osmophloeum* (found in Taiwan) inhibits planktonic cultures of many gram-positive and gram-negative bacteria, including MRSA (methicillin-resistant *Staphylococcus aureus*). Early studies described the major active compound in cinnamon oil as cinnamaldehyde. Cinnamaldehyde inhibits the proton motive force, respiratory chain, electron transfer, and substrate oxidation, disconnecting oxidative phosphorylation, inhibition of active transport, depletion of pool metabolites, and disruption of synthesis of RNA, DNA, lipids, polysaccharides, and proteins. Furthermore, volatile oils and their constituents exhibit hydrophobicity that allows them to partition into and disrupt the lipid bilayer of the cell membrane, so the membranes become more penetrable to protons. Large-scale leakage from bacterial cells or the loss of crucial molecules and ions eventually causes bacterial cell death. For example, exposure of S. *epidermidis* to cinnamon oil kills planktonic cells or staphylococci in biofilms.

Thus, cinnamon oil could be a valuable component in a combination antimicrobial drug therapy. According to the US FDA, “concurrent development of two or more novel drugs for use in combination generally will provide less information about the safety and effectiveness of the individual drugs than would be obtained if the individual drugs were developed alone.” Of course, this depends on many factors. The development stage of at which the separate drug components cease to be studied independently alters the amount of information available. FDA states that “because codevelopment will generally provide less information about the safety and effectiveness of the individual drugs, it will present greater risk compared to development of an individual drug.” However, cinnamon oil, as a substance already in the food supply, has the advantage of millions of person-years of exposure and safety data. The purpose of this study is to estimate the amount of exposure to cinnamon through selected foods in the United States in order to propose an initial target level for pharmacokinetic studies of a novel combination drug.

## Methods

### Assessment of Cinnamon Use

An assessment of the consumption of cinnamon by the U.S. population was conducted. Estimates for the intake of cinnamon were based on the approved food uses and maximum use level in conjunction with food consumption data included in the National Center for Health Statistics’ (NCHS) 2009-2010, 2011-2012, and 2013-2014 National Health and Nutrition Examination Surveys (NHANES) (CDC, 2006; USDA, 2012; Bodner-Montville et al, 2006). Calculations for the mean and 90th percentile intakes were executed for the representative food uses of cinnamon combined. The consumptions were reported for these <u>seven population groups:</u>

1. infants, age 0 to 1 year
2. toddlers, age 1 to 2 years
3. children, ages 2 to 5 years
4. children, ages 6 to 12 years
5. teenagers, ages 13 to 19 years
6. adults, ages 20 years and up
7. total population (all age groups combined, excluding ages 0-2 years)

## Food Consumption Survey Data

### NHANES Survey Description

The National Health and Nutrition Examination Survey (NHANES) is a biannually planned series of studies devised to measure the health and nutritional status of children and adults in the United States. The survey is distinctive in that it integrates interviews and physical examinations. NHANES is an important program operated by the National Center for Health Statistics (NCHS). The NHANES program was launched in the 1960s and has been orchestrated as a series of surveys paying particular attention to varying population groups or health topics. In 1999, the NHANES became a continuous program that has a dynamic focus on a diverse health and nutrition assessments to meet newly developing needs. The NHANES interview incorporates demographic, socioeconomic, dietary, and health-related questions. The examination section comprises medical, dental, and physiological assessments, as well as laboratory tests conducted by qualified medical personnel (CDC, 2018).

Discoveries made using NHANES are employed to calculate the prevalence of important diseases and risk factors for conditions. Data are used to measure nutritional status and its correlation with health encouragement and disease prevention endeavors. NHANES discoveries constitute the foundation for national standards for statistics such as height, weight, and blood pressure. Data from the NHANES are utilized in epidemiological research and health sciences studies that help develop solid public health plans, design and direct the health agenda and assistance available, and increase the health understanding of the US (CDC, 2018).

The most recent National Health and Nutrition Examination Surveys (NHANES) for the years 2013-2014 are available for public examination. NHANES are conducted as a continuous, annual survey, and are published online in 2-year cycles. In each cycle, approximately 10,000 people across the U.S. complete the health examination portion of the survey. Any combination of consecutive years of data collection is considered a nationally representative sample of the U.S. population. It is well known that the length of a dietary survey affects the estimated consumption of individual users and that short-term surveys, such as the typical 1-day dietary survey, overestimate consumption over longer time periods (Hayes et al, 1995). Because two 24-hour dietary recalls conducted on 2 non-consecutive days (Day 1 and Day 2) are available from the NHANES 2013-2014 surveys, these data were used to generate estimates for the current intake analysis.

The NHANES provide the most suitable data for evaluating food-use and food-consumption patterns in the United States, containing 2 years of data on individuals selected via stratified multistage probability sample of civilian non-institutionalized population of the U.S. NHANES survey data are collected from individuals and households via 24-hour dietary recalls administered on 2 non-consecutive days (Day 1 and Day 2) throughout all four seasons of the year. Day 1 data are collected in-person in the Mobile Examination Center (MEC), and Day 2 data are collected by telephone in the following 3 to 10 days, on different days of the week, to obtain the targeted degree of statistical independence. The data are obtained by first selecting Primary Sampling Units (PSUs), which are counties throughout the U.S. Small counties are aggregated to obtain a minimum population size. These PSUs are subdivided and households were chosen within each subdivision. One or more participants within a household are interviewed. Fifteen PSUs are visited each year. For example, in the 2009-2010 NHANES, there were 13,272 persons selected; of these 10,253 were considered respondents to the MEC examination and data collection. 9754 of the MEC respondents provided complete dietary intakes for Day 1 and of those providing the Day 1 data, 8,405 provided complete dietary intakes for Day 2. The release data does not necessarily incorporate all the questions asked in a section. Data items may have been removed due to confidentiality, quality, or other reasons. As a result, it is possible that a dataset does not completely match all the questions raised in a questionnaire section. Each data file is edited to include only those sample persons permitted for that particular section or component, so the numbers vary.

In addition to collecting data on the types and quantities of foods being ingested in the US, the NHANES surveys collect socioeconomic, physiological, and demographic information from individual participants in the survey, such as sex, age, height and weight, and other variables useful in characterizing consumption. The inclusion of this information allows for further assessment of food intake based on consumption by specific population groups of interest within the total population.

Sample weights were incorporated with NHANES surveys to compensate for the potential under-representation of intakes from specific population groups as a result of sample variability due to survey design, differential non-response rates, or other factors, such as deficiencies in the sampling frame (CDC, 2006; USDA, 2012).

#### Statistical Methods

Consumption data from individual dietary records, detailing food items ingested by each survey participant, were collated by computer in Matlab and used to generate estimates for the intake of cinnamon by the U.S. population. Estimates for the daily intake of cinnamon represent projected 2-day averages for each individual from Day 1 and Day 2 of NHANES data; these average amounts comprised the distribution from which mean and percentile intake estimates were produced. Mean and percentile estimates were generated incorporating sample weights in order to provide representative intakes for the entire U.S. population. “All-user” intake refers to the estimated intake of cinnamon by those individuals consuming food products containing cinnamon. Individuals were considered users if they consumed 1 or more food products containing cinnamon on either Day 1 or Day 2 of the survey.

### Food Usage

#### Food Data

Food codes representative of use were chosen from the Food and Nutrition Database for Dietary Studies (FNDDS) for the corresponding biennial NHANES survey (see Table 1). In FNDDS, the primary (usually generic) description of a given food is assigned a unique 8-digit food code (CDC, 2006; USDA, 2012).

**Table 1.** Selected Foodcodes Representative of Use. (Cinnamon concentrations, mean +/- SD)

| **Foodcode** | **Food Name** | **Concentration (mg/100 g)** |
| --- | --- | --- |
| 63101130 | Motts applesauce | 0.05 +/- 0.005 |
| 63101140 | Townhouse applesauce | 0.39+/- 0.09 |
| 53111000 | Gingerbread | 0.86 +/- 0.05 |
| 51160100 | Cinnamon honey buns | 8.4 +/- 0.2 |
| 51113010 | Cinnamon swirl bread | 31.1 +/- 2.8 |
| 57304100 | Cinnamon Life cereal | 1.8 +/- 0.2 |
| 57305165 | Cinnamon Grahams | 6.8 +/- 1.2 |
| 57103100 | Apple Cinnamon Cheerios | 12.3 +/- 0.6 |
| 57125000 | Cinnamon Toast Crunch | 18.7 +/- 1.4 |
| 53220030 | Apple cinnamon newtons | 0.04 +/- 0.01 |
| 92306800 | Chai Tea cookies | 2.7 +/- 0.1 |
| 53231400 | Whole wheat and cinnamon cookies | 15.8 +/- 1.3 |
| 13210110 | Bread pudding | 49 +/- 0.01 |
| 13210410 | Rice pudding | 1.9 +/- 0.3 |

## Results and Discussion

The mean mass of each population group retrieved from the NHANES 2013-2014 database appears in Figure 1. The group with the lowest average body mass is ages 0-1 and the group with the highest mass is ages 20+.

**Figure 1.**
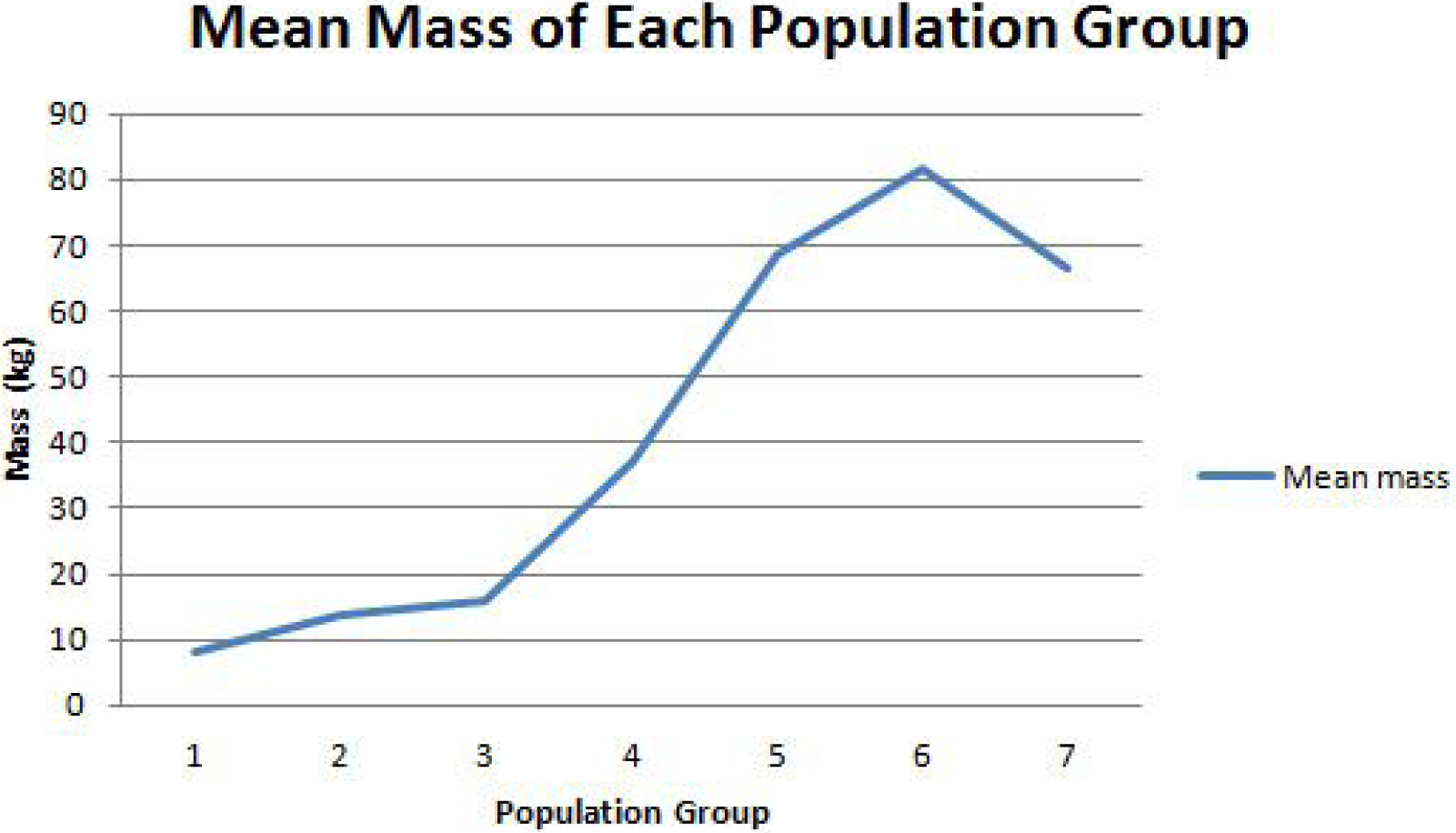
The highest average mass is the population group with ages 20 and up.

The percentage of users in each population group of the selected cinnamon-containing foods is depicted in Figure 2. The lowest consuming group was infants with ages 0-1, where no masses of consumption of the listed foods were reported. The highest consuming group was ages 6-12. In general, children were more frequent consumers of cinnamon-containing foods.

**Figure 2.**
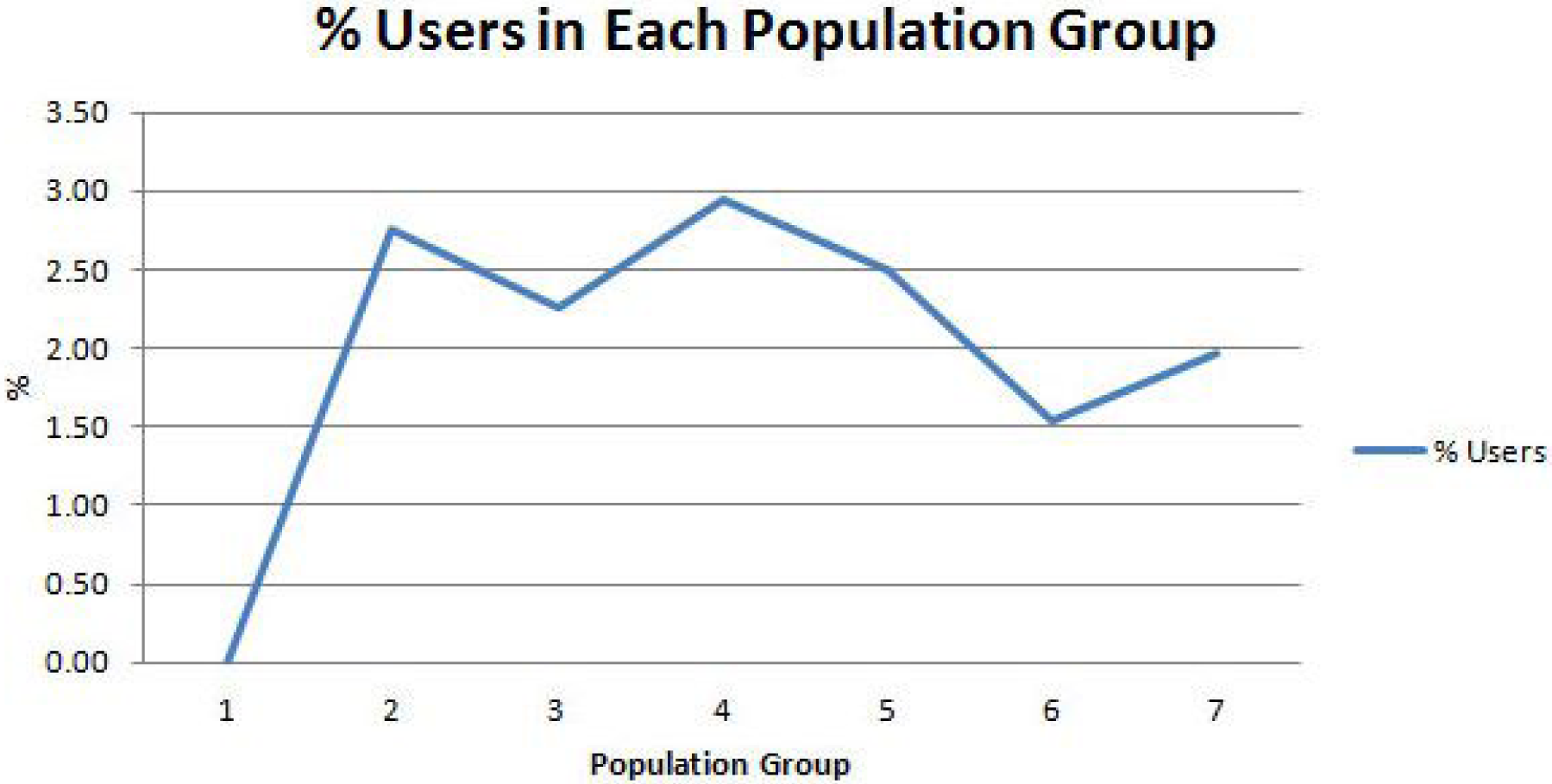
In general, children were more frequent consumers of the targeted cinnamon-containing foods than adults.

The expected daily intake of cinnamon in each population group from the targeted foods is shown in Figure 3. The heaviest consumers of the cinnamon-containing foods listed in Table 1 were teenagers from 13 to 19 years of age.

**Figure 3.**
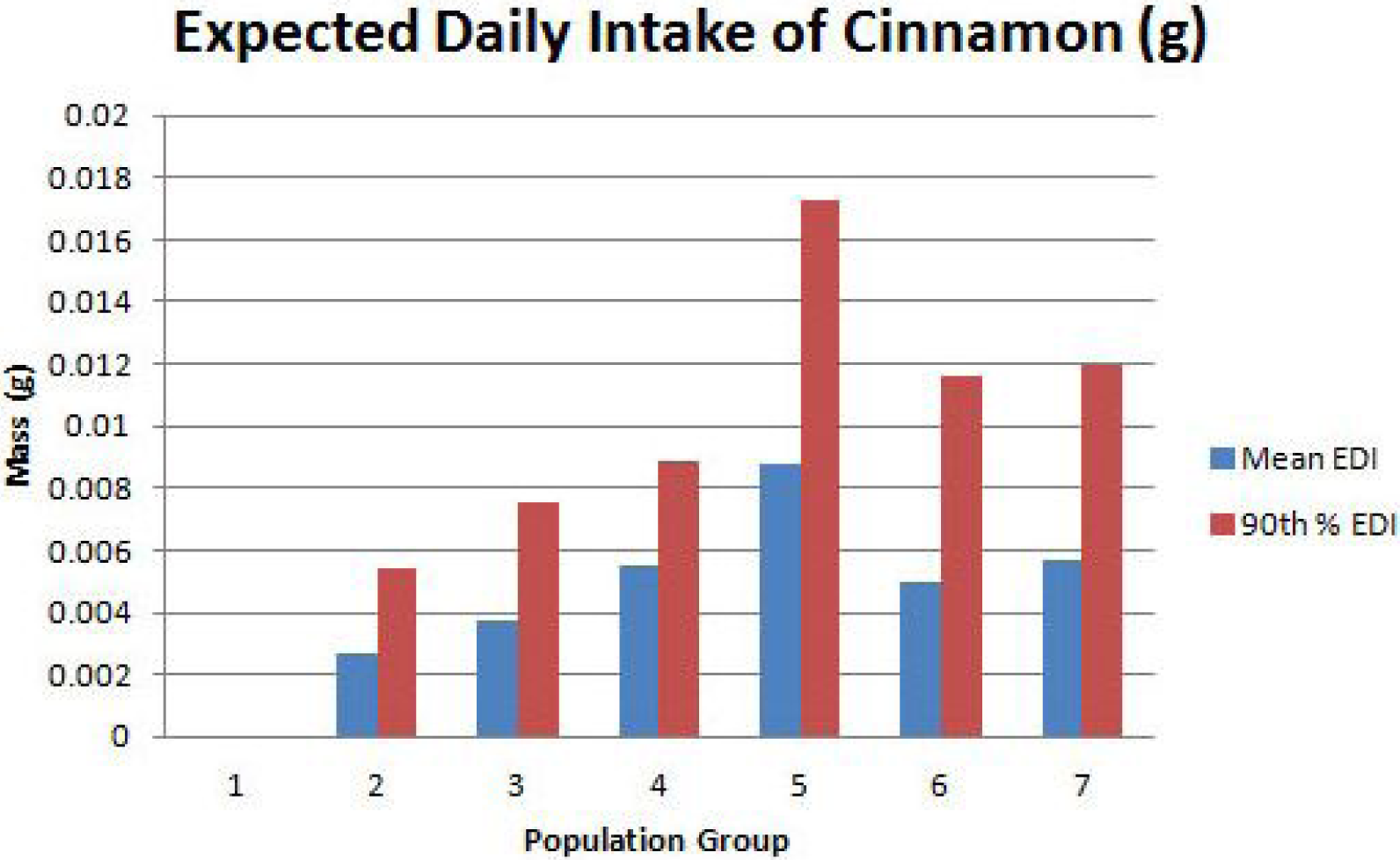
EDI of cinnamon from the targeted foods in Table 1. The heaviest consumers were in the 13-19 years of age group.

The weight-adjusted expected daily intake of cinnamon in each population group from the targeted foods is shown in Figure 4. The heaviest consumers of the cinnamon-containing foods listed in Table 1 on a weight-adjusted basis were children from 2 to 5 years of age.

**Figure 4.**
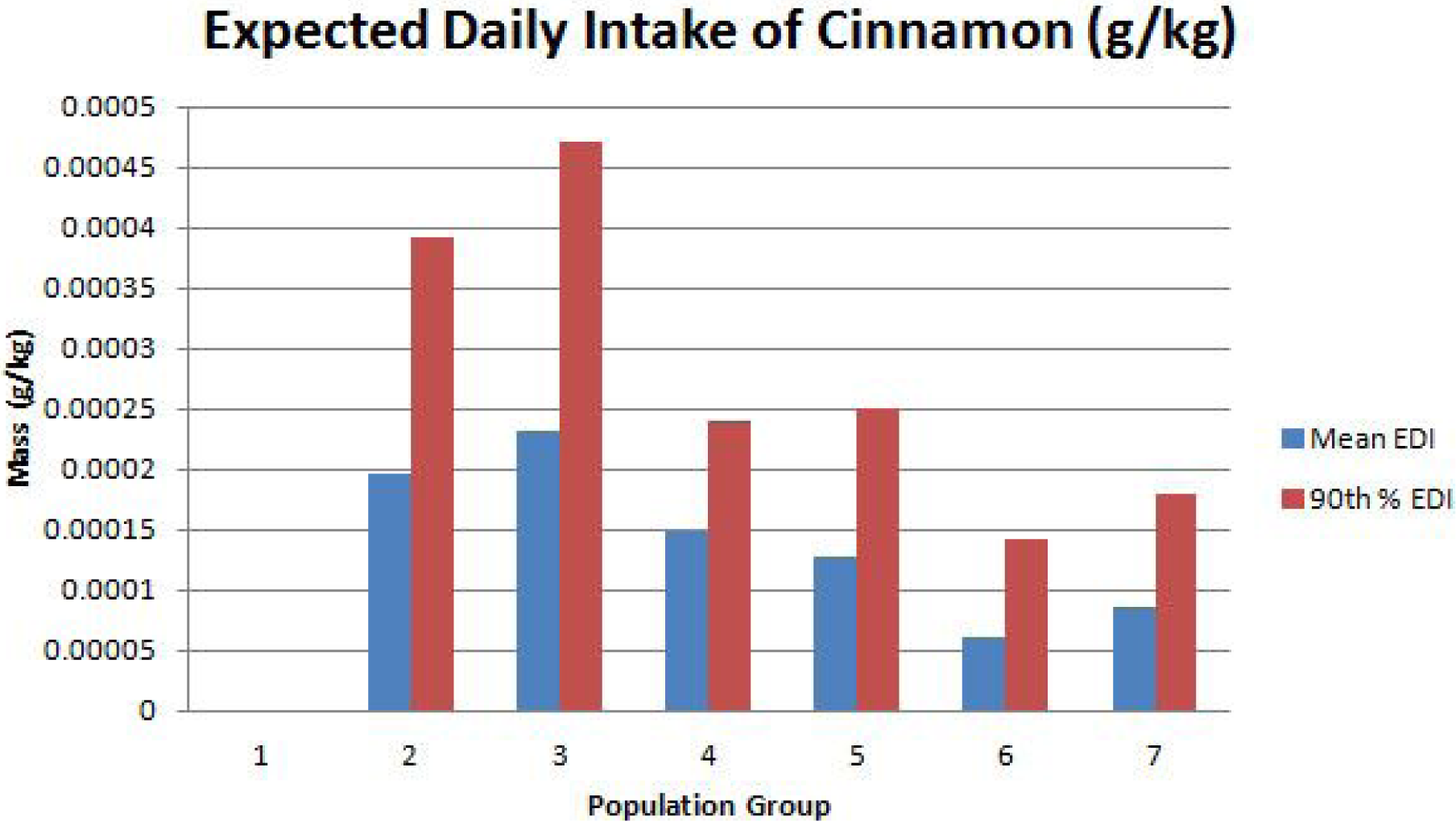
Weight-adjusted EDI of cinnamon from the targeted foods in Table 1. The heaviest consumers on a weight-adjusted basis were in the 2-5 years of age group.

## Conclusion

Biofilms protect bacteria from their environment, including drugs that may be added to their environment. Fifty percent or more of bacteria can exist in biofilms, which act like coral reefs to encapsulate and shield the cells. The most well-known biofilm may be dental plaque (Lu, 2007), but biofilms can also grow on medical devices like implants and catheters, where they cause potentially lethal infections. The cooperative advantages of growing in a biofilm enable bacteria to resist the activity of antibiotics far more effectively than free-swimming (planktonic) bacteria. In the struggle against biofilms, the challenge is to penetrate the matrix. Cinnamon can inhibit biofilm formation and disrupt existing biofilms, enabling an new antibiotic to work more effectively.

The mean daily exposure to cinnamon in the 20+ age group from the selected foods is 0.0050 g, and the 90th percentile exposure is 0.0116 g. On a weight-adjusted basis, the mean exposure to cinnamon in the 20+ age group from the selected foods is 0.0000617 g/kg body weight, and the 90th percentile exposure is 0.00014 g/kg body weight. A maximum dose of 0.00014 g/kg body weight is a reasonable dose starting point for a pharmacokinetic trial of a combination formulation containing cinnamon following washout.

## Credit

The project described was supported by the National Science Foundation through ACI-1053575 allocation number BIO170011, and the National Center for Research Resources and the National Center for Advancing Translational Sciences, National Institutes of Health, through Grant UL1TR001998. The content is solely the responsibility of the authors and does not necessarily represent the official views of the NIH.

